# Senescence and DNA Damage–Induced Inflammation Drive Heart Failure with Preserved Ejection Fraction in Cardiovascular Kidney Metabolic Syndrome

**DOI:** 10.64898/2026.04.13.718331

**Authors:** Dao-Fu Dai, Jun-yi Zhu, Meng Gao, Kaihao Wang, Nastaran Daneshgar, Xiaoping Yang, Virginia S. Hahn, Monica Vladut Talor, Daniela Cihakova, Avi Rosenberg, Antentor O. Hinton, Zhe Han

**Author notes:** Correspondence: Dao-Fu Dai, MD, PhD (lead contact), Cardiovascular and Renal Pathology Division, Department of Pathology, Johns Hopkins University School of Medicine, 632E Ross Building, 720 Rutland Ave, Baltimore, MD, 21205, or Zhe Han, PhD, University of Maryland School of Medicine, 670 West Baltimore Street, Baltimore, MD 21201, USA. equal contribution (Dao-Fu Dai, Jun-yi Zhu).

## Abstract

**Introduction:** Heart failure with preserved ejection fraction (HFpEF) is strongly associated with cardiometabolic comorbidities, including obesity, diabetes, hypertension, chronic kidney disease and aging, yet the mechanistic contribution of cellular senescence to HFpEF pathogenesis remains poorly defined.

**Methods and Results:** To model clinically relevant HFpEF, we subjected p16-3MR mice to a novel chronic “four-hit” cardiovascular-kidney-metabolic stress regimen (10 months of a high-fat diet, low-dose streptozotocin, L-NAME, and aging). These mice developed a robust HFpEF phenotype characterized by left ventricular hypertrophy, impaired diastolic function (reduced E′/A′ and elevated E/E′), preserved ejection fraction, reduced −dP/dt, exercise intolerance, pulmonary congestion, and increased cardiac CD68⁺ macrophage infiltration. Cardiac proteomics identified 821 proteins significantly altered by four-hit stress. Selective genetic ablation of p16⁺ senescent cells using ganciclovir ameliorated HFpEF phenotypes, reduced cardiac p16 expression and inflammation, and normalized proteomic remodeling, without affecting body weight or glycemic status. Comparative network analysis of mouse and human HFpEF cardiac proteomes revealed highly concordant upstream regulatory networks, prominently involving cell-cycle control, DNA damage responses, and inflammatory signaling. Immunohistochemical analysis of human HFpEF cardiac biopsies confirmed increased p16, γH2AX, STING, IRF3, NF-κB p65, and CD68⁺ macrophages, mirroring the murine findings. The 4-Hit mice also developed chronic diabetic kidney disease with increased kidney inflammation, both of which were attenuated by Senolytic therapy. Mechanistically, the cGAS-STING (cyclic GMP-AMP synthase – stimulator of interferon genes) is activated in response to damaged DNA, which in turn activates the downstream immune responses, including NF-κB and interferons. Cross-species validation further demonstrated that combined metabolic stress impaired cardiac function and nephrocyte function in *Drosophila.* Cardiac and nephrocyte dysfunctions were independently rescued by cardiomyocyte-specific and nephrocyte-specific inhibition of the cGAS-STING pathway, respectively. In human iPSC-derived cardiomyocytes, irradiation and palmitate induced senescence, DNA damage sensing via ZBP1, and activation of the cGAS-STING-IRF3 signaling axis; ZBP1 knockdown or senolytic treatment suppressed this inflammatory axis.

**Conclusions:** Across mouse, human, fly, and human iPSC models, our findings identify DNA damage–driven senescence and ZBP1–cGAS–STING signaling as conserved, causal mechanisms linking cardiovascular-kidney-metabolic comorbidities to HFpEF, highlighting senescence and innate immune pathways as promising therapeutic targets.

## Introduction

Approximately 50% of patients with HF have preserved ejection fraction (HFpEF), and this proportion rises in the elderly^1^. Despite recent positive clinical trials for HFpEF, these medications may be limited by eligibility and side effects, particularly in the elderly. HFpEF is increasingly known as a complex systemic disorder that develops with multiple comorbidities^2^, such as diabetes, hypertension, obesity, chronic kidney disease, and old age^3^. Inflammation has been implicated in HFpEF^4^. However, the mechanistic role of senescence and inflammaging in cardiometabolic syndrome, particularly in HFpEF, is not well understood.

Cellular senescence is a hallmark of aging that impairs tissue maintenance and repair. It is characterized by the arrest of cell division in mitotic cells, coupled with a senescence-associated secretory phenotype (SASP) that includes several proinflammatory and profibrotic cytokines. The current study examines the intricate relationship among senescence, the DNA damage response, and inflammation in HFpEF, which is driven by aging and cardiometabolic stress.

To elucidate the role of senescence, we applied a novel, clinically relevant multiple-hit cardiovascular-kidney-metabolic stress, including 10 months of high-fat diet (HFD), low-dose streptozocin, L-NAME, and middle age (designated as 4-Hit) in p16-3MR mice. The p-16-3MR reporter mouse model is a powerful tool for understanding the pathobiology of cellular senescence in aging and disease. The p16-3MR mice express a trimodality reporter fusion protein under the control of the p16^INK^ promoter. The 3MR contains Renilla luciferase, red fluorescent protein (RFP reporter), and truncated herpes simplex virus 1 thymidine kinase. Since most senescent cells express p16^INK4a^, these p16-3MR mice highlight p16-positive senescent cells that can be ablated by ganciclovir, thus creating a genetic senolytic therapy. In this study, we demonstrate that ablation of senescent cells alleviates HFpEF in the 4-Hit model. We show significant concordance between cardiac proteomic signatures in the 4-Hit mouse hearts and human HFpEF. In particular, we find consistent upregulation of pathways involved in senescence, DNA damage sensing and response, and pro-inflammatory pathways. Using a similar high-fat, high-sucrose feeding of the middle-aged *Drosophila* model, we demonstrate the mechanistic role of the pro-inflammatory cGAS (cyclic GMP-AMP synthase – stimulator of interferon genes) - STING (Stimulator of Interferon Genes) pathway. We then validate the relevance of this mechanism to human HFpEF using human induced pluripotent stem cells (iPS-CM) and human HFpEF cardiac biopsies.

## Method (Further details in Supplemental File)

### Mice and Cardiac Studies

All animal procedures were performed in accordance with the Institutional Animal Care and Use Committee at our institution (M023M236) and the guidelines from the NIH Guide for the Care and Use of Laboratory Animals. The high fat diet has 60 % kcal from fat (D12494) and control diet has 10 kcal% (D12450J). Both were ordered from Research Diets.Inc. Because previous studies have established the model of HFpEF in male mice^5^, we used male p16-3MR mice aged 3-4 months with a C57BL/6J background in this study.

Using Vevo 2100 VisualSonics (Toronto, ON, Canada), echocardiography was performed in conscious, gently manually restrained mice injected with 0.1 mg midazolam. The images were obtained using a 30-MHz linear array transducer. The analysis was done blinded to the treatment group, using the biplane area–length method for LV mass and ejection fraction. Diastolic function was measured using tissue Doppler imaging of the mitral annulus (E’, A’) and conventional mitral inflow (E).

### Quantitative proteomics analysis by Tandem Mass Tag Method

Frozen cardiac apical tissues were lysed in RIPA buffer. Cardiac total protein lysates were reduced, alkylated, and purified by chloroform/ methanol extraction before digestion with trypsin. The peptide mix was labeled using a Tandem Mass Tag 11-plex isobaric label reagent set (Thermo). We used an Orbitrap Eclipse Tribrid mass spectrometer (Thermo) to acquire the data.

Following data acquisition and database search, the MS3 reporter ion intensities were normalized using ProteiNorm^6^. The data was normalized using VSN^7^ and analyzed using ProteoViz to perform statistical analysis using Linear Models for Microarray Data (limma) with empirical Bayes (eBayes) smoothing to the standard errors.

### Human HFpEF biopsy bioinformatics analysis and Immunohistochemistry (IHC)

We performed bioinformatics analysis by eXpression2Kinase analysis (https://maayanlab.cloud/X2K/; Icahn School of Medicine, Mount Sinai, New York)^8^ using the list of significantly altered proteins from two current mouse heart datasets (4-Hit/control and 4-Hit+Senolytic / 4-Hit), compared with the HFpEF proteome (vs. control) previously reported in our paper^9^ (Supplementary Tables). The previous HFpEF proteomics study used label-free Data-Independent Acquisition (DIA-MS). We focused on HFpEF2 subjects as they had higher body mass index and greater cardiac proteomic changes (n=11).

### Immunohistochemistry and immunoblotting

Multiple sections of 11 confirmed HFpEF cases were compared with 5 controls from autopsy hearts of decedents without significant cardiovascular diseases. Ventana Discovery Ultra autostainer (Roche Diagnostics) was used for CD68, p16 and γH2AX. Due to limited availability of samples, serial IHC was performed manually for NF-κBp65, IRF3, and STING. Quantitative analysis of the CD68+ cell count, p16+ cell count, or integrated densities of other IHC staining was performed by a pathologist (blinded to the treatment group), using HALO (Area Quantification FL Algorithm, Indica Laboratory, Albuquerque, NM) or ImageJ, as described^10, 11^. The antibody information is listed in Table 1.

**Table 1.** Antibodies.

| **Antibodies** | **Dilution ratio** | **Catalog / Lot #** | **Company** |
| --- | --- | --- | --- |
| *Immunostaining* |  |  |  |
| CD68 | 5000 | m0814/20014287 | DAKO |
| γH2AX | 1000 | ab81299 | Abcam |
| IRF3 | 600 | MA532348 | Thermo Fisher |
| NF-κB p65 | 500 | #8242 | Cell Signaling Technology |
| p16 | 1000 | 705-4793 | Ventana |
| STING | 100 | #13647S | Cell Signaling Technology |
| *Immunoblot* |  |  |  |
| β-actin | 500 | sc-47778 | Santa Cruz Biotechnology |
| cGAS | 1000 | 29958-1-AP | Proteintech |
| GAPDH | 500 | sc-32233 | Santa Cruz Biotechnology |
| p-IRF3 (Ser396) | 1000 | #29047 | Cell Signaling Technology |
| IRF3 | 2000 | #80519-1-RR | Proteintech |
| p-STINGser366-d7c3s | 1000 | #19781 | Cell Signaling Technology |
| STING | 2000 | 19851-1-AP | Proteintech |
| ZBP1 | 1000 | A13899 | Abclonal |

### Sucrose and fat feeding in *Drosophila*

Flies were maintained on standard food (Meidi LLC) at 25°C under standard conditions. The following *Drosophila* lines were obtained from the Bloomington Drosophila Stock Center (BDSC): UAS-*CG7194*-RNAi (ID_41855), UAS-*Sting*-RNAi (ID_31565), and *w^1118^* (ID_3605). The *Hand*-Gal4 driver was generated in our lab and used to express RNAi-based silencing (-IR) constructs in the heart. *Hand*>*w^1118^* was used as a control. The cardiac marker R94C02::tdTomato was provided by Rolf Bodmer (Sanford Burnham Research Institute, CA).

The 10% Sucrose (Sigma-Aldrich) and/or 2% fat (Coconut oil) were dissolved in water and added to standard fly food. Under normal conditions, water alone was added to the standard food. Standard fly food was obtained from Meidi (V100) and is based on the BDSC cornmeal food recipe by the Bloomington Drosophila Stock Center. Flies were kept at 25°C for normal and sucrose and/or fat treatment.

### Live imaging of adult *Drosophila* heart

Cardiac function in adult *Drosophila* was measured using ZEISS SteREO Discovery.V12 with a 305-color camera (ZEISS, Jena, Germany). Young (4-day-old) or middle-age (21-day-old) adult female flies were used. Female flies have larger body size that facilitates dissection. Flies were dissected in artificial hemolymph [70 mmol/l NaCl (Carolina), 5 mmol/l KCl, 1.5 mmol/l CaCl2·2H_2_O, 4 mmol/l MgCl_2_, 10 mmol/l NaHCO_3_, 5 mmol/l trehalose, 115 mmol/l sucrose, and 5 mmol/l HEPES in H_2_O. All chemicals were purchased from Sigma-Aldrich. Each fly was gently placed on a plate with petroleum jelly (Vaseline) for immobilization, with the ventral aspect facing the microscopy source. For each genotype, 10 flies per genotype were used. ZEISS SteREO Discovery.V12 was used to record the adult heart rhythm and heart wall movement in the same position, i.e., the cardiac chamber in the abdominal segment A3 of each fly. Each measurement was obtained in 5 distinct positions within each abdominal segment (A2, A3, and A4), and these were averaged to obtain the cardiac diameter for that fly. ImageJ software (version 1.49) was used to process the images, blinded to the treatment group. The diastolic dimension and systolic diameter were processed, measured, and determined based on three consecutive heartbeats. The heart rate was determined by counting the total number of beats that occurred during a 30-second recording.

### The 10 kD dextran uptake assay and *Drosophila* Nephrocyte size

Dextran uptake by nephrocytes was assessed *ex vivo* in young (4-day-old) or middle-age (21-day-old) female adult flies to expose nephrocytes in artificial hemolymph. Nephrocytes were dissected from young (4-day-old) or middle-age (21-day-old) female adult flies and kept in artificial hemolymph. Nephrocyte size was determined using the area measurement function in ImageJ (version 1.52a). Confocal images were obtained using a ZEISS LSM900 microscope and ZEN Blue (edition 3.0) acquisition software.

### Differentiation of induced pluripotent stem cell-derived cardiomyocytes, siRNA transfection and treatment with Senolytic agents

Human induced pluripotent stem cells (iPS cells) from normal young adult (ATCC ACS-1026) were grown on plates coated with Matrigel (Corning Life Sciences) and fed with 50% SFM XF (ATCC ACS-3001) / 50% mTeSR Plus (STEMCELL Technology). Cardiomyocyte differentiation was performed using a well-established protocol^12^. Tri-iodo-L-thyronine (T3; 100 nmol/L) and dexamethasone (1 μmol/L) were added from day 14 to promote iPS-CM maturation. Approximately 25 days after differentiation, iPS-CM were subjected to 10 Gy of X-ray, followed by 10 days of palmitate (100 μM). Eight days after palmitate, 15 nM of ZBP1 siRNA (siRNA ID.s37486, ThermoFisher) was used to transfect iPSC-CMs, diluted to 100 nM in RPMI-1640 and Lipofectamine 2000 (Invitrogen). The iPS-CM was transfected for 48 hours and used for subsequent experiments. For a separate group receiving senolytic agents, Dasatinib (20μM) and Quercetin (15μM) were added for 48 hours. Ten days after palmitate and IR, all cells were lysed in RIPA buffer containing protease and phosphatase inhibitors.

### Statistical Analyses

Statistical analyses were performed using GraphPad Prism 9 or Stata 10. Continuous variables were represented as mean ± standard error. Shapiro-Wilk normality tests were applied to define violation of normality. For data that does not violate normality, we applied ANOVA for evaluations involving multiple groups, followed by Sidak post hoc tests for pairwise comparisons. Student’s t-test (unequal variance) was applied for two group comparisons. Skewed data were assessed with non-parametric methods: Kruskal–Wallis test (for multiple groups). Nonparametric Dunn’s tests were used for two groups to adjust for multiple comparisons. Each figure legend outlines the statistical methods. The p<0.05 was considered statistically significant. The limma statistical method was employed for -omics data analysis, with appropriate adjustments made for multiple comparisons to control for potential false discovery rates (FDR:1% by Benjamini-Hochberg method).

## Results

### Cardiometabolic stress model (4-Hit) induces senescence and heart failure with preserved ejection fraction (HFpEF)

HFpEF is well-known to be associated with multiple comorbidities, including diabetes, hypertension, obesity, old age, and chronic kidney disease. To model HFpEF patients with comorbidities, we applied a novel “four-hit” cardiovascular-kidney-metabolic (CKM) stress (abbreviated 4-Hit thereafter), which consist of: 1) long-term high fat diet (60 % kcal from fat) for ∼10 months to induce obesity as a metabolic stress, 2) low-dose streptozocin (STZ) injection (40mg/kg/day, up to 5 days) to induce diabetes and hyperglycemia, 3) L-NAME (N^[w]^-nitro-l-arginine methyl ester, 0.5 g/L) added to the drinking water to induce endothelial dysfunction and 4) middle age (13-14 month-old). The experimental design is summarized in Figure 1A. We used the p16-3MR (hereafter p16) mouse model to highlight cells expressing p16 senescence marker (RFP reporter), which enables a selective ablation or clearance of these p16+ senescent cells by ganciclovir (GCV). At the end of the experiment, the 4-Hit mouse hearts demonstrate accumulation of RFP-positive senescent cells, with RFP intensity increased by >2-fold compared with those in mice fed with a control diet. Three cycles of the senolytic therapy GCV successfully decreased the senescent cells (Figure 1B) and significantly decreased the RFP signal of senescence markers (Figure 1C).

**Figure. 1.**
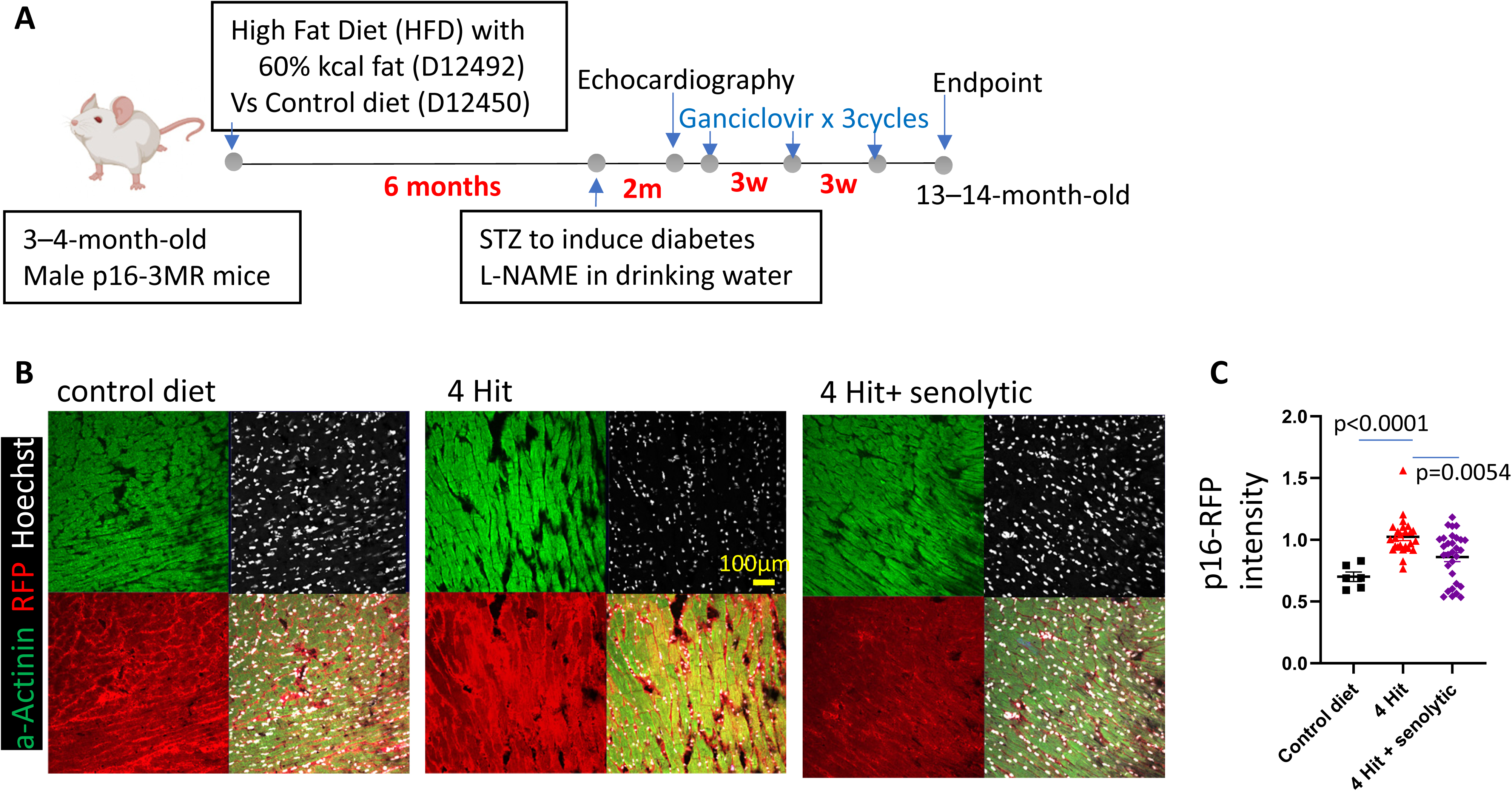
Experimental design and p16 Signals. (A) The young p16-3MR of C57Bl6/J mice were subjected to 4-Hit cardiometabolic stress, including 10 months of high-fat diet (60% kcal fat), streptozocin (STZ), L-NAME, and middle-age. 3 doses of Ganciclovir did genetic ablation in p16-3MR mice. (B). Representative immunofluorescent images of p16 (red) and α-actinin (green) staining of left ventricular sections from each experimental group. (C). Relative intensity of red (p16) signals. N=6-12 mice; significance by non-parametric tests.

The 4-Hit CKM mice became obese after 6 months of high fat diet and developed hyperglycemia after low-dose STZ (Figure S1A). The hyperglycemia was not affected by senolytic therapy (Figure S1B). At the end of the experiment, the 4-Hit mice showed significant increases in body weight and heart weight-to-tibia length (Figures 2A and 2 B), indicating obesity and cardiac hypertrophy. There was significantly increased lung weight/tibia length in 4-hit mice (Figure.2C), suggesting lung congestion related to heart failure. Pathological examinations revealed that 4-Hit mice had myocardial inflammation, predominantly composed of macrophages (Figure 2D, confirmed by positive CD68 staining, Figure S2). Echocardiographic evaluation of 4-Hit mice revealed a significant increase in LV mass (Figure 2G). The left ventricular ejection fraction (LVEF) was preserved (Figure 2H). The diastolic function in 4-Hit mice was significantly impaired, measured by abnormal Tissue Doppler È/À (Figure.2I) and E/E’ (Figure 2J), which suggest impaired relaxation and increased LV filling pressure. We performed a Millar catheter measurement of intracardiac pressure to complement the echocardiographic data. The 4-Hit mice displayed a mild, ∼16 % decrease in +dP/dt (the rate of LV pressure change during systole), and a substantial decrease in -dP/dt (the rate of LV pressure change during diastole) by ∼40%, suggestive of a much stronger impairment of diastolic function. The 4-Hit mice undergoing the Treadmill exercise test had more than 50% decrease in exercise tolerance. Altogether, these findings confirm the development of the HFpEF in the 4-Hit CKM mice.

**Figure. 2.**
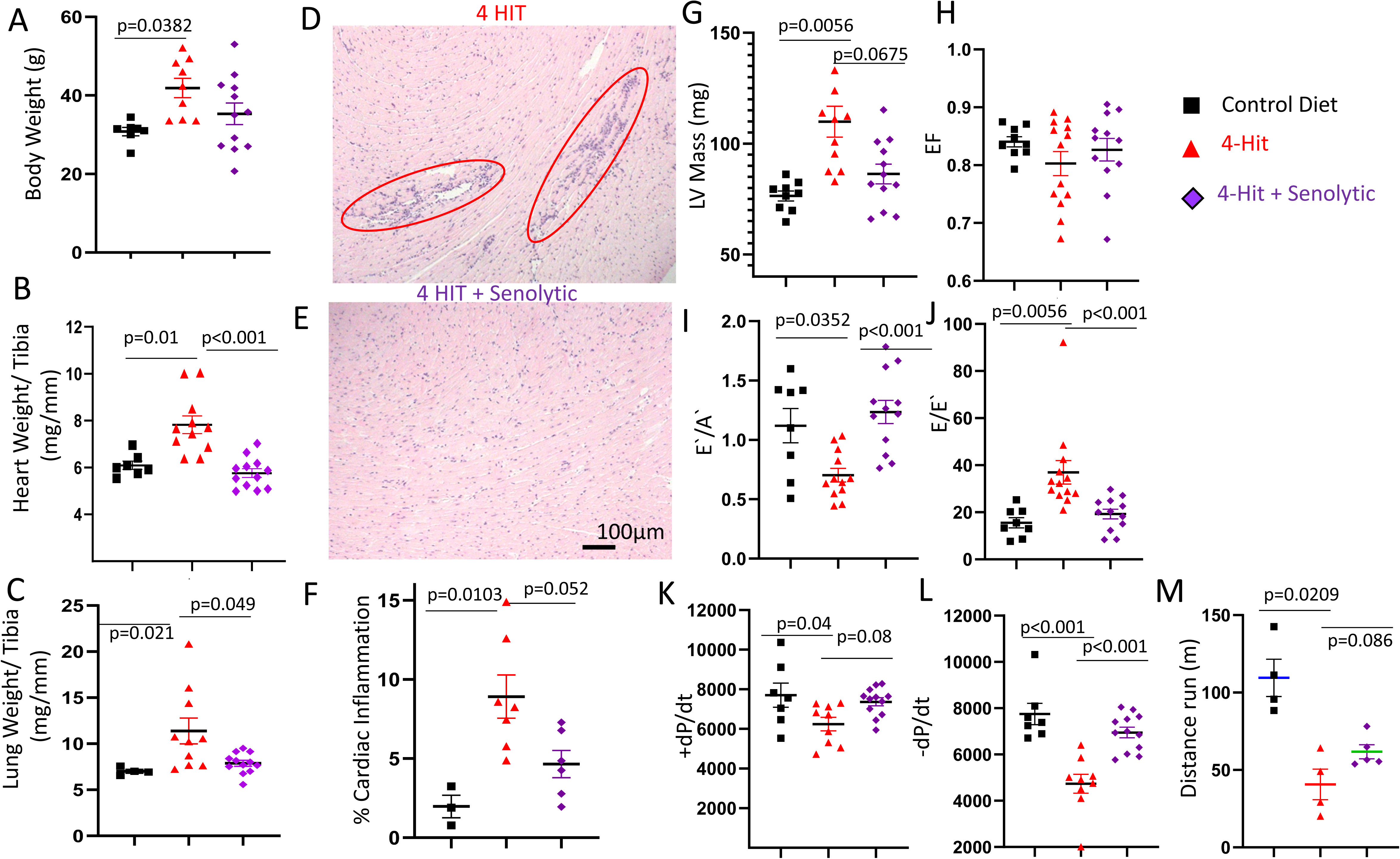
Cardiometabolic syndrome (4-Hit) and HFpEF in mice. Three experimental groups of p16-3MR mice: control diet only, 4-Hit, and 4-Hit treated with Senolytic (ganciclovir). (A) Body weight (g), (B) Heart weight (normalized to tibia length), Lung weight (normalized to tibia length).(D-E). Representative H&E-stained myocardial tissues with mononuclear cells inflammatory infiltrates in 4-Hit and 4-Hit treated with Senolytic. Echocardiography (G-J) showed increased LV mass (G), no significant change in EF % (H), diastolic dysfunction shown by (I) E’/A’ and (J) E/E’. By Millar catheter, (K) intracardiac pressure changes over time (K) +dP/dt during systole and (L) –dP/dt during diastole. By Treadmill exercise, (M) distance run (m). N=3-12. Statistical tests by ANOVA followed by Sidak post-hoc tests (A,F,K,L), or non-parametric Kruskal-Wallis and Dunn’s tests (B,C,F,G,I,J,M).

### Genetic ablation of senescent cells (senolytic therapy) improved HFpEF

Senolytic therapy with three cycles of GCV did not significantly affect blood glucose (Figure S1) or body weight (Figure.2A), but significantly attenuated cardiac hypertrophy, lung congestion, and myocardial inflammation (Figure.2B-F).

Echocardiography showed that 4-Hit mice treated with GCV had significantly lower LV mass and significant improvement in diastolic function (È/À and E/È), without changing the LVEF (Figure.2G-J). Millar catheter showed that senolytic mildly improved +dP/dt (p=0.08) and significantly restored -dP/dt diastolic function, comparable to the control diet group (Figure.2K-L). There was also a mild improvement in Treadmill exercise tolerance (p=0.07, Figure.2M). These findings suggest that senolytic therapy ameliorates the HFpEF induced by CKM stress.

### Genetic senolytic therapy ameliorated diabetic kidney disease and the associated inflammation

Low-dose STZ induced diabetes in obese mice (Figure S1A). These 4-Hit mice developed many pathologic features of chronic diabetic kidney disease, including mesangial expansion (Figure 3A-B), thickening of glomerular basement membrane (Figure 3D) and tubular basement membrane (Figure 3E), as well as tubular atrophy and interstitial fibrosis (% of Trichrome blue, Figure 3C). The 4-Hit kidneys also display clusters of mononuclear inflammatory infiltrates (Figure S3A), which are associated with chronic injury in diabetic kidney disease^13^. Immunohistochemistry for CD68 reveals increased macrophage infiltration, both within the glomeruli and in the interstitium (Figure S3B). The 4-Hit CKM stress also induced upregulation of IL-6, a pro-inflammatory cytokine (also a component of SASP), and Caspase-1, an inflammatory protease within the inflammasome (Figure S3C-D). These findings suggest that CKM stress induced systemic multi-organ inflammation. Senolytic therapy significantly attenuated all of the inflammatory signals in the kidneys (Figure S3) and ameliorated the pathologic changes of diabetic kidney disease (Figure 3).

**Figure. 3.**
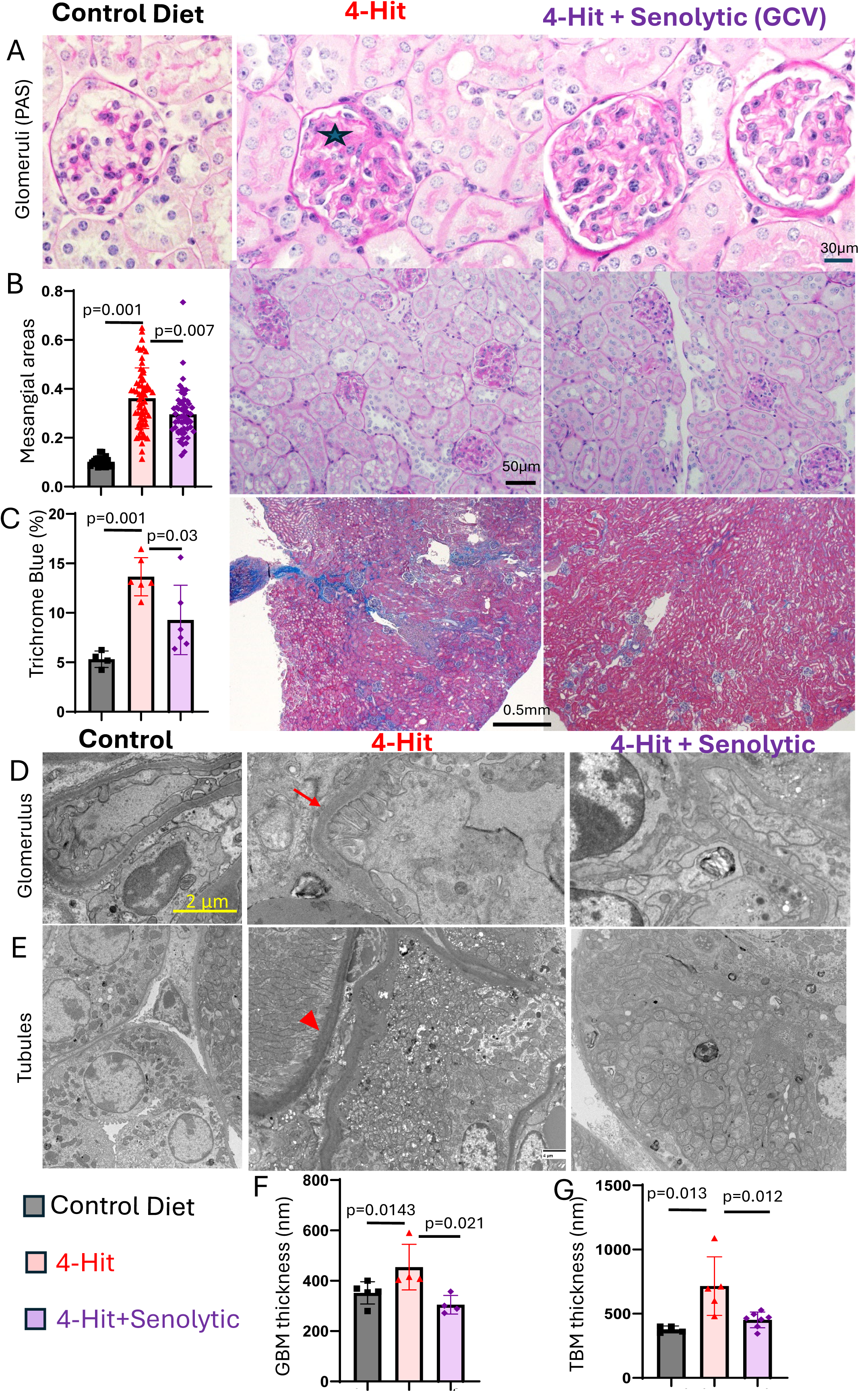
Kidney pathology in 4-Hit mice. Three experimental groups of p16-3MR mice: control diet only, 4-Hit, and 4-Hit treated with Senolytic (ganciclovir). (A) Representative high magnification images of glomeruli (PAS staining) showing expanded mesangial areas (*) in 4-Hit, (B) Mesangial areas / total glomerular areas and representative PAS images, (C) Trichrome blue (%) and representative images, (D) Representative Electron Microscope (EM) images showing thickening of Glomerular basement membrane (GBM, arrow), (E) Representative EM images showing thickening of Tubular Basement Membrane (TBM, arrowhead), (F) GBM thickness (nm), (G) TBM thickness (nm); N=4-6. Statistical tests by ANOVA followed by Sidak post-tests (B-C) or non-parametric tests (F-G).

### Mechanistic Insights from Mouse and Human Quantitative Cardiac Proteomics

To gain mechanistic insights into the molecular signaling underlying 4-Hit CKM stress induced HFpEF and how these were ameliorated by senolytic treatment, we performed quantitative proteomics using the Tandem Mass Tag method, comparing the relative abundance of the whole cardiac proteome in mice receiving a control diet, 4-Hit, and 4-Hit + senolytic treatment. We analyzed the ratio (log fold change) of the same proteins in 4-Hit / control (4-Hit effect) and separately for 4-Hit+senolytic / 4-Hit (senolytic treatment effect). Out of 3525 proteins identified in cardiac lysate of all mouse heart samples (Supplementary Table), there were 821 proteins significantly altered by 4-Hit, and 320 proteins significantly changed in the senolytic vs untreated 4-Hit group. Both 4-Hit effect and senolytic effect were visualized using volcano plots (Figures S4A and S4B).

We performed bioinformatic analysis using eXpression2Kinases Web (X2K, Mount Sinai Bioinformatics) to link proteome changes to the upstream cell signaling networks^8^. X2K Web infers upstream regulatory networks from signatures of significantly altered (differentially expressed) proteins. By combining transcription factor enrichment analysis, protein-protein interaction network expansion, and kinase enrichment analysis, X2K Web produces inferred networks of transcription factors, proteins, and kinases that are predicted to regulate the set of significantly altered proteins. We performed X2K analysis using the current mouse heart proteome dataset (821 for 4-Hit vs control; 320 for senolytic effect on 4-Hit) and compared to the HFpEF patients’ heart DIA proteomics that was reported in our recent paper (1294 proteins significantly altered in obese HFpEF2 vs control)^9^.

The predicted upstream regulatory networks in X2K were presented as the top 20 results from Transcription Factor enrichment and Kinase enrichment analyses. Compared with the human HFpEF proteome (Figure.4A-C), the 4-Hit-mouse heart proteome (Figure.4D-F) displays highly overlapping upstream transcription factors and kinases (highlighted in Figure 4D, E). The top overlapping transcription factors include TAF1, MYC, YY1, E2F1, MAX, ATF2, NFYB, BRCA1, etc. The top overlapping upstream kinases include cyclin-dependent kinases (CDK1, CDK4, CDK2, CDC2), several Mitogen-Activated Protein Kinases: MAPK14 (p38), MAPK1 (ERK2), MAPK3 (ERK1), MAPK8 (JNK1), and several kinases in DNA damage and repair pathways (ATM, DNAPK, PRKDC, and ABL1). In addition to the overlapping transcription factors with the other two groups, the Senolytic-treated 4-Hit mouse hearts (right column of Figure 4) also show Interferon Regulatory Factor 3 (IRF3), RELA (an active component of NF-κB), and STAT3, a critical regulator of immune response. Overall, the eXpression2Kinases Network in both human HFpEF and 4-Hit mouse hearts emphasizes the regulation of cell cycle, DNA damage, and DNA damage response/repair, as well as inflammatory pathways. The effect of Senolytic therapy highlights many components of inflammatory pathways (IRF3, RELA, STAT3, JNK, etc.), suggesting crucial roles for inflammatory and DNA damage response pathways as potential underlying mechanisms of the benefit of Senolytic therapy.

**Figure 4.**
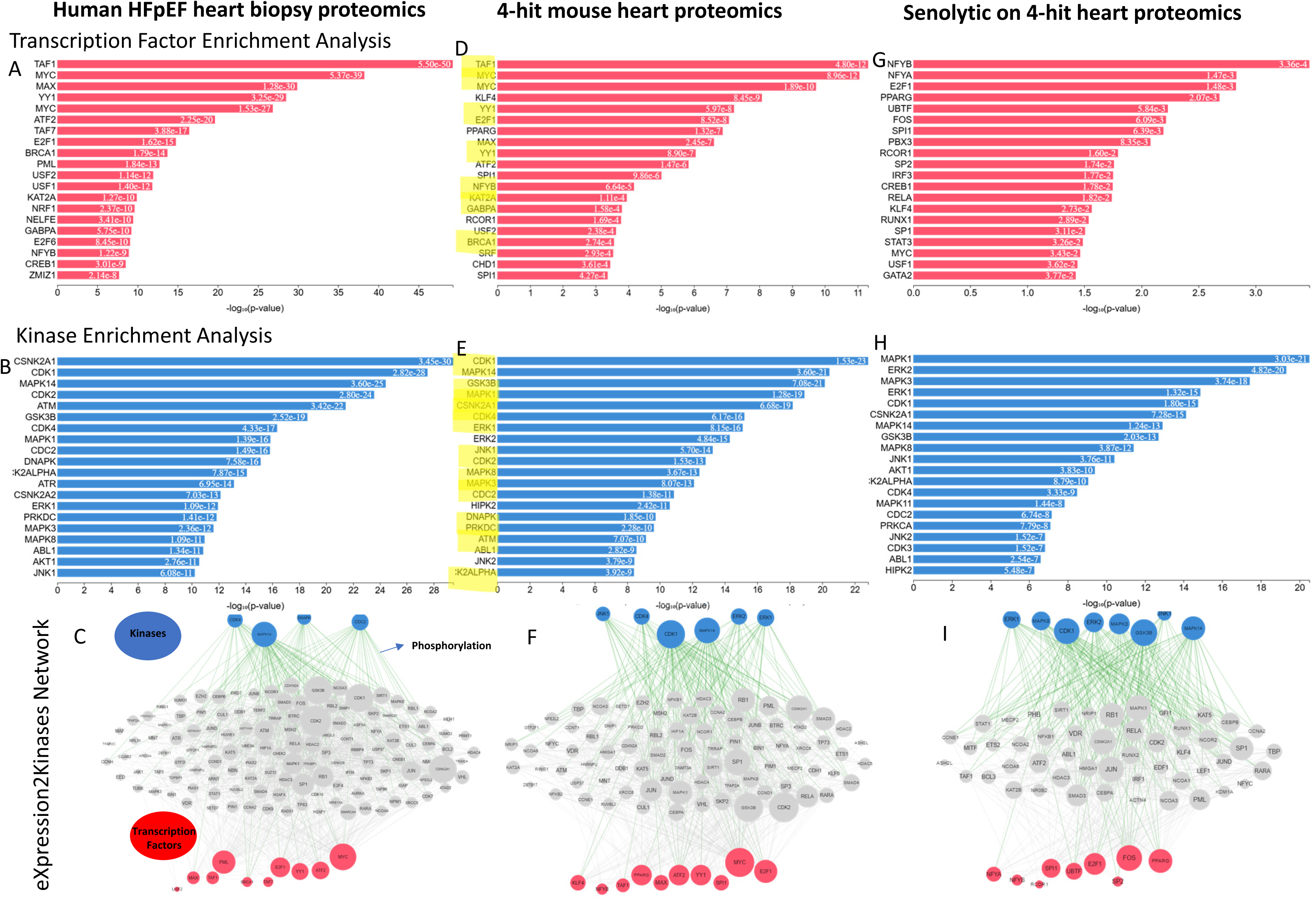
Top 20 Upstream Transcription Factors and Kinases Network. (A) Top 20 transcription factors inferred from 1294 proteins significantly altered in human HFpEF heart biopsy proteome (vs. normal control), (B). Top 20 kinases and (C) Network analysis inferred from the same dataset. (D) Top 20 transcription factors inferred from 821 proteins significantly altered in 4-hit mouse heart proteome (vs. control), (E). Top 20 kinases and (F) Network analysis inferred from the same dataset. (G). Top 20 transcription factors inferred from 320 proteins significantly altered in Senolytic + 4-hit mouse heart proteome (vs. 4-hit), (H). Top 20 kinases and (I) Network analysis inferred from the same dataset. Bioinformatic analysis by eXpression2Kinases. Yellow highlights overlap between 4-hit mouse heart proteome and human HFpEF cardiac proteome.

### Increased senescence and inflammation in human HFpEF hearts

To validate the clinical importance of DNA damage response and inflammatory pathways in human HFpEF, we performed immunohistochemistry of HFpEF endomyocardial biopsy samples matching the proteomics samples^9^ and the eXpression2Kinases analysis. Compared to controls, myocardial biopsies from HFpEF patients show significantly higher CD68+ macrophage infiltration (Figure 5A)^10^, stronger signals of p16 senescence marker (Figure 5B) and γH2Ax (Figure 5C), a marker of oxidative DNA damage. cGAS (cyclic GMP-AMP synthase) is activated by cytosolic (damaged) DNA, which in turn activates STING (stimulator of interferon genes), triggering downstream immune responses, including NF-κB and interferon. The HFpEF myocardium displayed significant and concordant increases in the signals of STING (Figure 5D), IRF3 (Figure 5E), and p65/RelA (component of NF-κB, Figure 5F).

**Figure 5.**
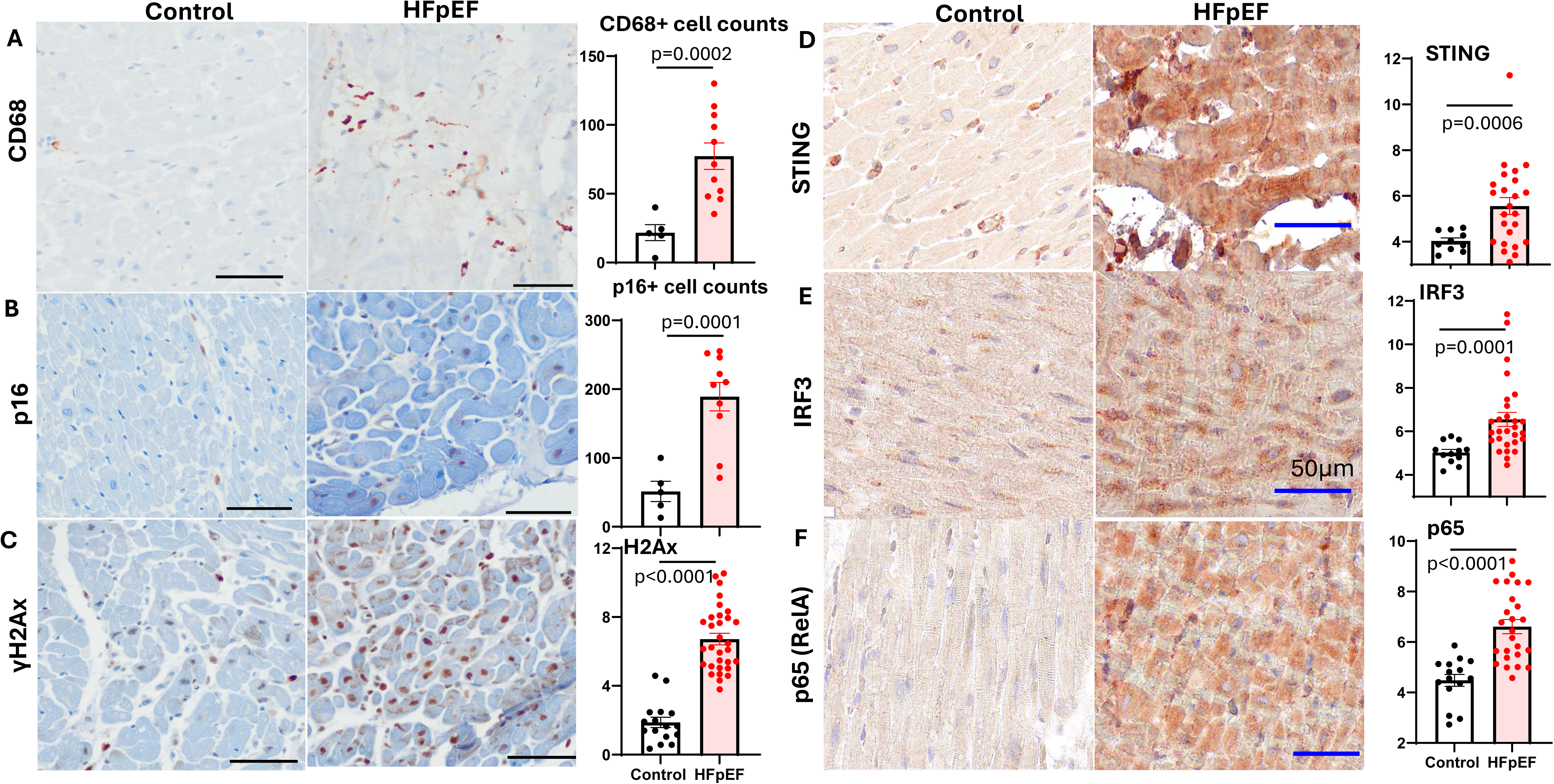
Immunohistochemistry of human HFpEF vs control heart sections. Representative images and quantitative analysis of (A) CD68+ cells /mm^2^, (B) p16+ cells /mm^2^, (C). γH2Ax (integrated density), (D).STING (integrated density), (E). IRF3 (integrated density), (F). P65 (RelA) (integrated density). Scale bars: 50µm. N=5-11. Significance determined by Student’s t-tests.

### Multiple-hit modeling of high dietary sucrose and fat in middle-aged *Drosophila* caused cardiac functional decline

To model multiple hit cardiometabolic stress, we fed the flies with 10% sucrose and/or 2% fat, then examined cardiac function using a quantitative video imaging-based method^14^. As shown in Figure 6, there was no significant difference between young and middle-aged *Drosophila* cardiac function. Feeding either high sucrose or high fat in middle-aged flies led to increased diastolic and systolic diameters (Figure 6A, B, and C) and decreased fractional shortening / cardiac contractility (Figure 6D) as well as slower heart rate (Figure 6E). Consistent with this, critical indicators of cardiac function, such as stroke volume and cardiac output, were also reduced in response to high-sucrose or high-fat feeding (Figures 6F and 6 G). Notably, a combination of high-sucrose and high-fat feeding in middle-aged flies further potentiated cardiac dysfunction, which was more severe than either stress alone (Figure 6A-G). These findings reinforce the synergistic suppression of fly cardiac function by multiple hits of cardiometabolic stress.

**Figure 6.**
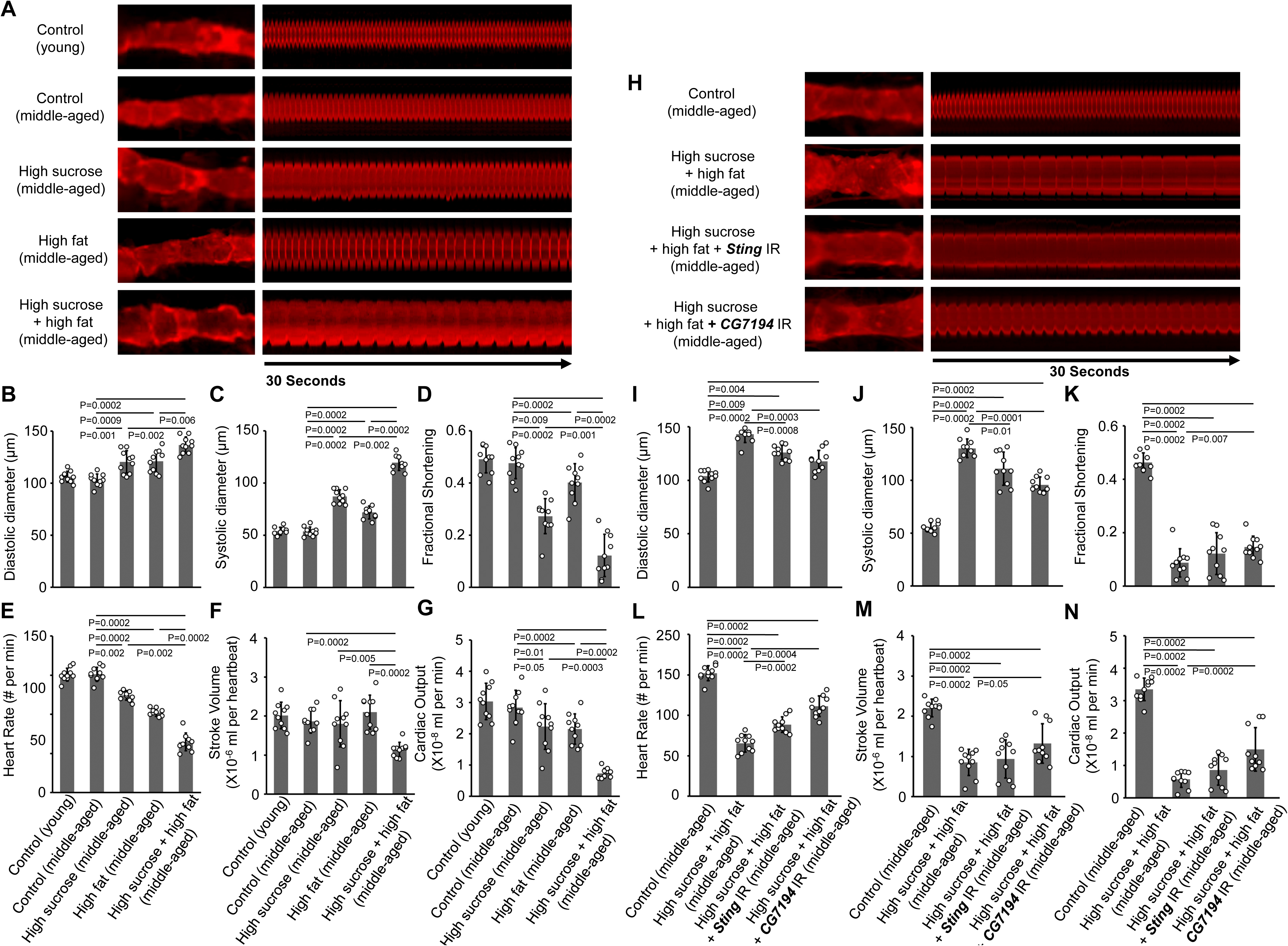
Inhibition of cGAS-STING pathway attenuated cardiac dysfunction in middle-aged *Drosophila* fed with high-sucrose and high-fat. **(A)** Images captured from heartbeat videos of young (4-day-old females) or middle-age (21-day-old females) *Drosophila*. Representative images show changes in cardiac function in flies with a normal diet (normal sucrose and fat) and those treated with a sucrose and/or fat diet. **(B)** Quantitation of adult *Drosophila* heart systolic diameter. n = 10 flies per genotype. **(C)** Quantitation of adult *Drosophila* heart diastolic diameter. n = 10 flies per genotype. **(D)** Quantitation of adult *Drosophila* heart rate. n = 10 flies per genotype. **(E)** Quantitation of adult *Drosophila* heart fractional shortening. n = 10 flies per genotype. **(F)** Quantitation of adult *Drosophila* heart stroke volume. n = 10 flies per genotype. **(G)** Quantitation of adult *Drosophila* heart cardiac output. n = 10 flies per genotype. **(H)** Images captured from heartbeat videos of middle-age (21-day-old females) *Drosophila* on a normal diet (normal sucrose and fat) and those fed with a high sucrose and high fat diet. Cardiac-specific driver *Hand*-Gal4 was used to knockdown the *CG7194* and *Sting*, the *Drosophila* homolog of the mammalian cGAS-STING pathway, in adult *Drosophila*. **(I)** Quantitation of adult *Drosophila* heart systolic diameter. n = 10 flies per genotype. **(J)** Quantitation of adult *Drosophila* heart diastolic diameter. N = 10 flies per genotype. **(K)** Quantitation of adult *Drosophila* heart rate. n = 10 flies per genotype. **(L)** Quantitation of adult *Drosophila* heart fractional shortening. n = 10 flies per genotype. **(M)** Quantitation of adult *Drosophila* heart stroke volume. n = 10 flies per genotype. **(N)** Quantitation of adult *Drosophila* heart cardiac output. n = 10 flies per genotype. Values were presented as means ± standard deviation (s.d). Kruskal–Wallis H-test followed by a Dunn’s test.

### The cGAS-STING pathway inhibition attenuated multiple-hit-induced *Drosophila* cardiac dysfunction

To elucidate the mechanisms, we examined whether genetic modification of the cGAS-STING pathway in the *Drosophila* heart could attenuate cardiac functional defects caused by combined high-sucrose/high-fat feeding. To achieve this, we combined the cardiac-specific driver *Hand*-Gal4^15^ with RNAi knockdown (UAS-*CG7146*-RNAi or UAS-*Sting*-RNAi) targeting CG7194 or Sting, the *Drosophila* homolog of mammalian *cGAS* and *STING*, respectively. To minimize the potential effects of the cGAS-STING pathway on *Drosophila* heart development, embryos from the progeny were initially maintained at a lower temperature (18°C), which suppresses GAL4 promoter activity and prevents premature knockdown of target genes^16^. The progeny resulting from crosses between *Hand*-Gal4 and UAS lines carried a cardiac-specific driver (*Hand*-Gal4) to induce RNAi-mediated silencing of either *CG7194* or *Sting*. After the appropriate developmental stage, these flies were shifted to a higher temperature (25°C) to enhance GAL4 activity, thereby activating the cardiac-specific knockdown of the cGAS-STING pathway. Simultaneously, flies were exposed to the combined high-sucrose/high-fat feeding. Under these conditions, silencing *CG7194* or *Sting* significantly alleviated the decline in cardiac function observed in multiple-hit middle-aged flies (Figure 6H-N), suggesting a critical role of cGAS/STING pathway in response to multiple-hit cardiometabolic stress.

### The cGAS-STING pathway inhibition attenuated nephrocyte dysfunction and shrinkage caused by a high-sucrose, high-fat diet in middle-aged *Drosophila*

Nephrocytes are podocyte-like cells that filter and endocytose to detoxify the unwanted substances from the *Drosophila* hemolymph^17, 18^. To determine the effect of multiple-hit stress on nephrocyte function, we used an *ex vivo* assay that measured the capacity of dissected nephrocytes to filter and endocytose the10 kDa fluorescent dextran particles. There was no significant difference in nephrocyte uptake function between young and middle-aged *Drosophila* fed a control diet (Figure 7A, B). However, we observed a significant reduction in dextran intensity in nephrocytes from middle-aged flies treated with high sucrose or high fat compared to those fed a normal diet.

**Figure 7.**
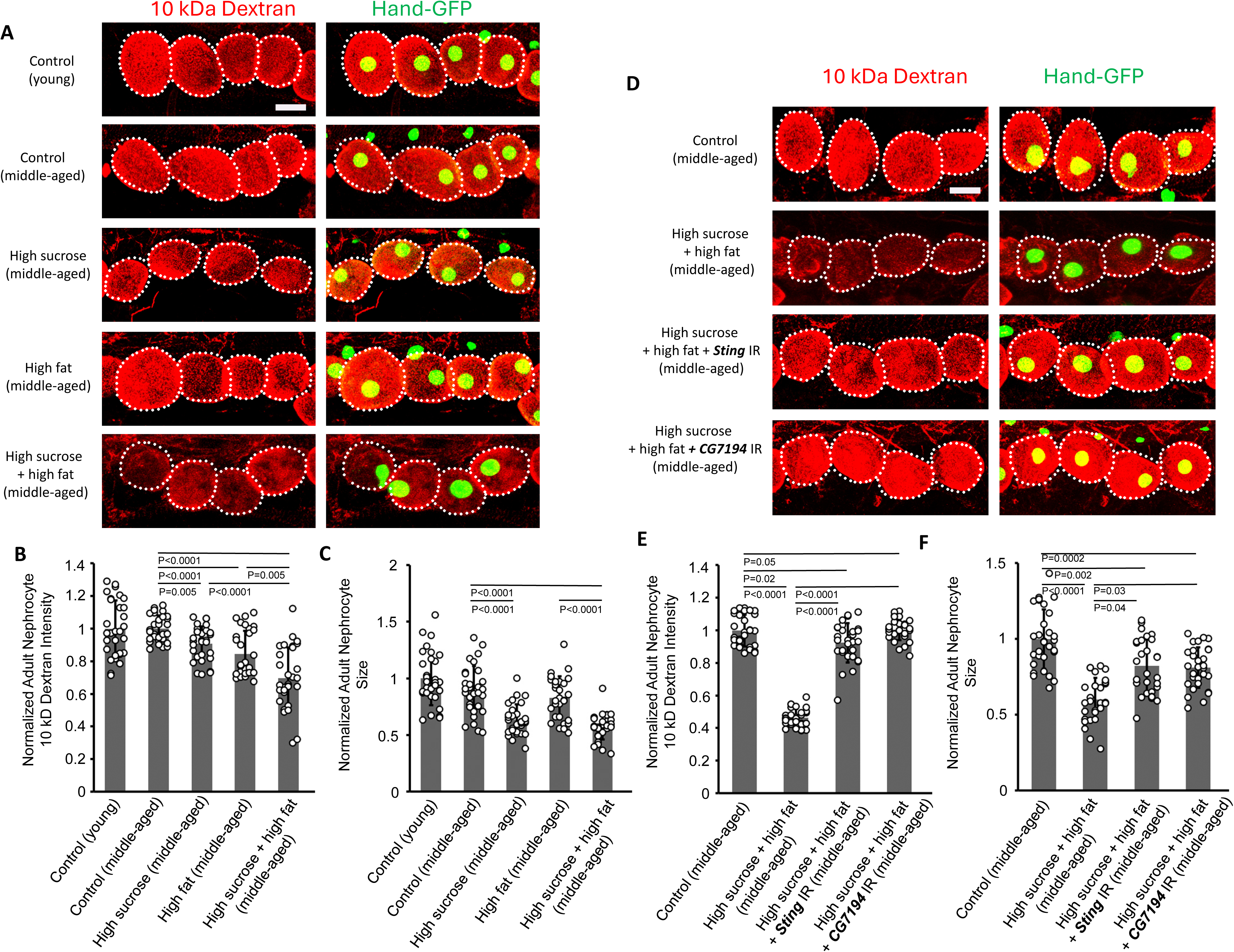
Inhibition of cGAS-STING pathway attenuated nephrocyte dysfunction in middle-aged *Drosophila* fed with high-sucrose and high-fat. **(A)** The 10 kDa dextran uptake (red) by nephrocytes from young (4-day-old adult females) and middle-aged (21-day-old adult females) flies on a control diet (normal sucrose and fat) and those fed with high-sucrose and/or high-fat diet. Hand-GFP transgene expression was visualized as green fluorescence concentrated in the nephrocyte nuclei (Hand-GFP; *Dot*-Gal4 flies). Scale bars: 15 µm. **(B)** Quantitation of 10kD dextran uptake, relative to uptake in young control flies (4-day-old adult females). n=30 nephrocytes from six flies per group. **(C)** Quantitation of adult nephrocyte size, relative to size in young control flies (4-day-old adult females). n=30 nephrocytes from six flies per group. The 10 kDa dextran nephrocytes uptake from middle-aged flies on a control diet, on a high-sucrose + high-fat diet, and the effect of cGAS-STING inhibition. Nephrocyte-specific driver *Dot*-Gal4 was used to knockdown the *CG7194* and *Sting*, the *Drosophila* homolog of mammalian cGAS-STING pathway, in adult Drosophila. Hand-GFP *transgene* expression was visualized as green fluorescence concentrated in the nephrocyte nuclei (Hand-GFP; *Dot*-Gal4 flies). Scale bars: 15 µm. **(E)** Quantitation of 10kD dextran uptake, relative to uptake in middle-aged control flies (21-day-old adult females). n=30 nephrocytes from six flies per group. **(F)** Quantitation of adult nephrocyte size, relative to size in middle-aged control flies (21-day-old adult females). n=30 nephrocytes from six flies per group. Results have been presented as mean ± s.d., normalized to the mid-aged control flies (21-day-old adult females). Kruskal–Wallis H-test followed by a Dunn’s test.

Notably, combined high-sucrose / high-fat significantly aggravated nephrocyte dysfunction, when compared with either (Figure.7A, B). In addition, nephrocyte size was significantly reduced following high-sucrose or high-fat feeding, and the combination induced more severe nephrocyte shrinkage compared with either stress alone (Figure 7A, C).

Next, we examined whether genetic modification of the cGAS-STING pathway in *Drosophila* nephrocytes could attenuate the nephrocyte functional defects caused by multiple-hits. To achieve this, we applied the nephrocyte-specific driver *Dot*-Gal4 with RNAi knockdown (UAS-*CG7146*-RNAi or UAS-*Sting*-RNAi) of *CG7194* or *Sting*, bypassing the developmental stage as mentioned above. Silencing *CG7194* or *Sting* significantly alleviated nephrocyte functional decline and cell shrinkage in middle-aged *Drosophila* subjected to high-fat and high-sucrose stress (Figure 7D-F).

### The cGAS/STING and ZBP1 are conserved in human senescent cardiomyocytes induced by irradiation and palmitate

To test if these pathways are relevant to human cardiomyocyte senescence, we differentiated cardiomyocytes from human induced pluripotent stem cells (iPSC-CMs). To induce senescence *in vitro,* we subjected iPSC-CMs to either ionizing radiation (IR) (10GY) or palmitate (100μM). Global transcriptomics analysis of iPSC-CM populations by RNA-sequencing revealed 1548 differentially expressed genes between IR-iPSC-CM and control iPSC-CM, and 201 differentially expressed genes between Palmitate-treated iPSC-CM and control iPSC-CM. We identified upregulation of several markers of cellular senescence that overlap with aging markers, including Growth differentiation factor 15 (GDF15) and CDKN1A (p16). Several CM proliferation markers, such as CDK1, CDC25, AURKB, as well as genes responsible for the condensation and stabilization of chromosomes during mitosis and meiosis (e.g. NCAPG) were downregulated in IR-iPSC-CM compared with controls (Figure 8A). Quantitative PCR of senescence markers showed significantly increased p16, p21, and IL1 in IR-treated iPSC-CM (Figure 8C), while palmitate treatment only showed a modest increase in these markers. IR-treated iPSC-CMs also display increased senescence-associated β-galactosidase (Figure S5B).

**Figure 8.**
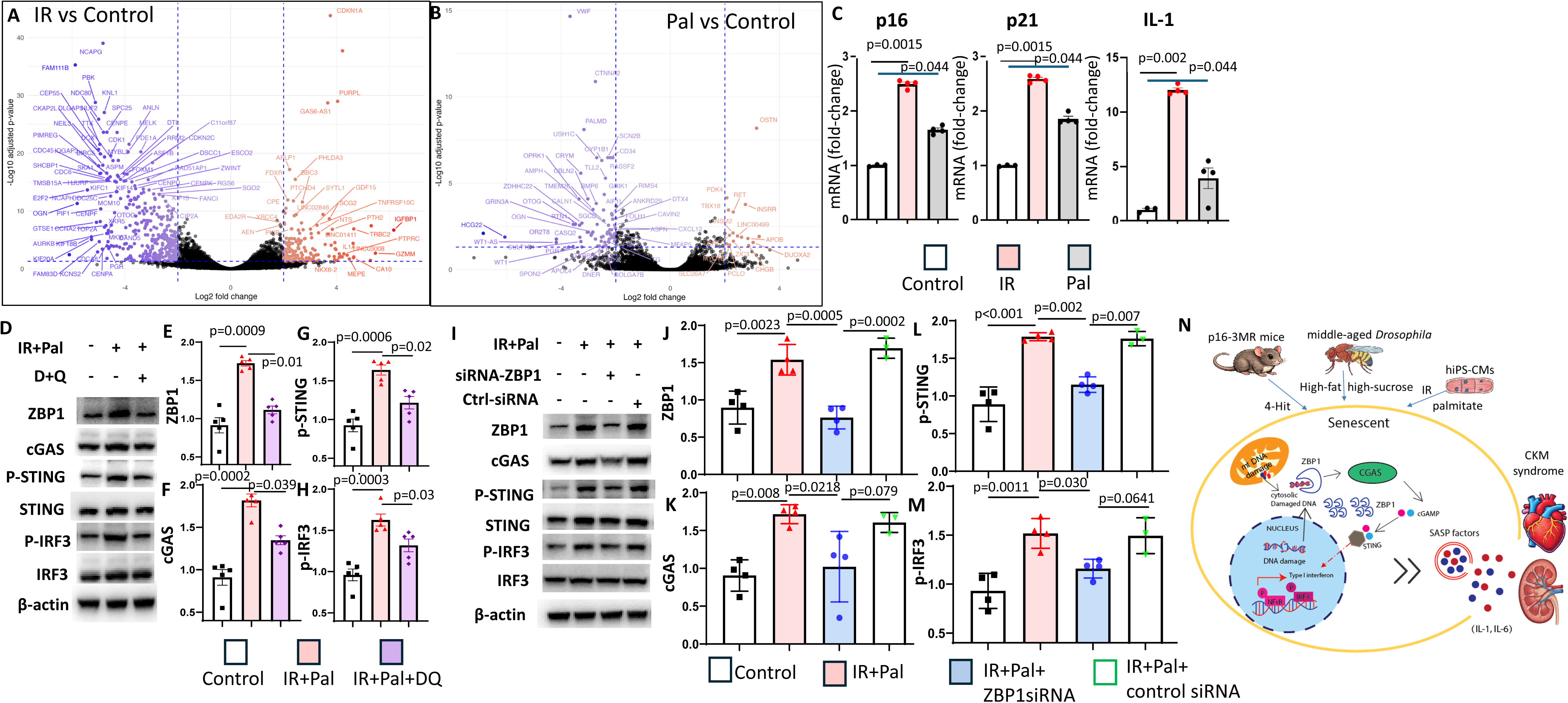
The human iPS-CM senescence. The volcano-plots of RNA-seq after (A) X-Ray irradiation (IR) or (B) Palmitate. (C). Quantitative PCR of p16, p21 and IL1. (D). Western blots of IR and Palmitate stress CM treated with D+Q and the quantitative analysis of (E). ZBP1, (F). cGAS, (G) phospho-/total STING, (H) phospho-/total IRF3. (I). Representative Western blots of IR and Palmitate stress CM treated with ZBP1 siRNA or control scrambled RNA and the quantitative analysis of (J).ZBP1, (K).cGAS, (L). phospho-/total STING and (M). phospho-/total IRF3. n=3-5, tested by non-parametric Kruskal-Wallis and Dunn’s tests. (N). Schematic diagram of signaling mechanisms.

We combined both IR and palmitate to model senescence and high fatty acid stress (similar to other models). In the stressed iPS-CM (Figure 8D), there were significant increases in both Z-DNA binding protein 1 (ZBP1, Figure 8E) and cGAS (Figure 8F). As cGAS activates STING, which activates/ phosphorylates IRF3 and p65/RelA, a component of NFkB, we also observed increased phosphorylation of both STING (Figure 8G) and IRF3 (Figure 8H). Treatment with senolytic agents (Dasatinib and Quercetin) suppressed ZBP1, which also attenuated cGAS, phosphorylation of STING, and phosphorylation of IRF3 (Figure 8D-H). Inhibition of ZBP1 by siRNA also attenuated cGAS/p-STING and p-IRF3 (Figure 8I-M). Taken together, these findings support a critical role for DNA damage induced-NF-κB and interferon pathways in response to senescence stressors, and senolytic agents significantly mitigated these signals. Consistently, the effect of genetic senolytic therapy was confirmed in 4-Hit CKM stressed mouse hearts. The 4-Hit significantly activated ZBP1/cGAS/STING pathways, all of which were mitigated by senolytic therapy (ZBP1, p=0.068; cGAS, p=0.058; STING phosphorylation, p=0.025, Figure S6).

## Discussion

Inflammation has been implicated in heart failure with preserved ejection fraction (HFpEF). However, the roles of senescence and inflammatory signaling mechanisms in HFpEF remain poorly defined. Our current study provides robust evidence for the mechanistic roles of senescence in HFpEF, as genetic senolytic therapy with ganciclovir in p16-3MR mice significantly ameliorated HFpEF. We further showed in middle-aged *Drosophila* that inhibition of cGAS/STING attenuated cardiac dysfunction induced by a high-sucrose, high-fat diet. Using iPS-CM stressed with X-ray irradiation and palmitate, we successfully modeled senescence. These senescent cardiomyocytes induced activation of ZBP1 and cGAS (both are sensors of cytosolic/damaged DNA) and downstream signaling through STING/IRF3 and NF-κB pathways. Treatment with the senolytic agents Dasatinib and Quercetin attenuated DNA damage sensors and downstream pathways, consistent with the findings in 4-Hit mice. We further confirmed that human HFpEF cardiac biopsies display a concordant activation of senescence, DNA damage, cGAS/STING, and inflammatory pathways. As summarized in the schematic diagram (Figure 8N), CKM stress leads to persistent DNA damage, which activates the ZBP1 and cGAS/STING pathway, amplifies NF-kB and interferon signals, induces p16+ senescence, and ultimately drives diastolic dysfunction and HFpEF. Removing senescent cells may attenuate the propagation of these signals and improve both HFpEF and diabetic kidney disease. Taken together, we elucidate the critical roles of senescence and DNA damage pathways in HFpEF using multiple models and validate these in human HFpEF.

HFpEF has become increasingly important as its prevalence is rising due to aging populations and increased rates of comorbidities such as obesity, hypertension, and diabetes^19^. The mortality and morbidity rates are marginally better than those with reduced ejection fraction (HFrEF)^20^. With limited effective treatment and frequent hospitalization, HFpEF imposes a major burden on healthcare and highlights a critical gap in medical research. Mice are the most extensively used pre-clinical models. The HFpEF features of diastolic dysfunction and preserved ejection fraction have been reported in aging^21^, angiotensin II infusion^22^, and many other rodent models of chronic hypertension^23^, either genetic^24^ or high-fat diet-induced obesity^25^. However, each model resembles some aspects of HFpEF but fails to meet the clinical criteria for HFpEF. The complex pathophysiology and multi-organ involvement in HFpEF make it challenging to model^26^. Recent studies suggest that HFpEF is a systemic disease closely associated with a cluster of cardiovascular stress, kidney dysfunction, and metabolic syndrome. Withaar et al reviewed the reported pre-clinical models and evaluated them against the clinical HFpEF score^26^. Among the well-accepted models are high-fat diet with nitric oxide synthase inhibitor (e.g., L-NAME) ^5^ and aging, high-fat diet, and angiotensin II infusion^27^. In this study, we extended the high-fat diet to 10 months to include an aging component and add a low-dose streptozocin to induce diabetes in these obese mice. This novel 4-hit CKM stress model closely recapitulates human HFpEF, particularly those with higher body mass index^9^. In addition, these mice also developed CKD due to diabetic nephropathy, compatible with systemic inflammation and multi-organ involvement within the cluster of cardiovascular-kidney-metabolic (CKM) syndrome.

The proposed signaling mechanism is summarized in Figure 8N. Cellular senescence is one of the most significant hallmarks of aging. Various cardiometabolic stresses can also induce it. The two most important characteristics of senescence are cell cycle arrest and the senescence-associated secretory phenotype (SASP). The DNA-damage response pathway (e.g. DNA double-strand breaks) induced upregulation of cyclin inhibitors, which is also observed in the aged heart^28^. The main SASPs include secretion of several proinflammatory cytokines (e.g., IL-1, IL-6), chemokines, and matrix metalloproteinases. These molecules may induce senescence of neighboring cells via paracrine signaling. The molecular link between senescence and chronic inflammation involves the cGAS/STING pathway in the ageing brain^29^. In response to chronic stress, damaged DNA from nuclei and mitochondria (mtDNA) is released into the cytoplasm, where it is recognized by the enzyme cGAS^30^. Once activated, cGAS produces cGAMP, a second messenger that binds to STING, triggering a downstream immune response, including the pro-inflammatory Senescence-Associated Secretory Phenotype (SASP)^30^. ZBP1 (Z-DNA binding protein 1) is another sensor that binds to Z-form nucleic acids, such as telomeric-repeat-containing RNA^31^ or damaged mtDNA^32^. Recent research suggests that ZBP1 is a critical amplifier within this axis, as shown in autoimmune photosensitivity^33^. The initial cGAS-STING signaling upregulates ZBP1, and then it becomes fully activated by sensing this non-canonical left-handed Z-DNA, such as from mtDNA, undergoing torsional stress^34^. This was observed in doxorubicin-induced cardiotoxicity^32^. The cooperative sensing mechanism stabilizes pro-inflammatory signaling complexes (e.g., RIPK1 and RIPK3), thereby promoting a robust and persistent inflammatory state. Overall, these contribute to the systemic “inflammaging” characteristic of biological aging^32^. Our current data highlight the critical role of these integrated inflammaging pathways in chronic CKM syndrome, including HFpEF and diabetic kidney disease. We further showed that the activation of immunomodulatory transcription factors NFkB and IRF3, together with the involvement of macrophages, may regulate inflammation and senescence through feedback loops in HFpEF hearts.

Our data provides robust evidence for the mechanistic role of senescence in HFpEF and diabetic kidney disease under cardiometabolic stress. This may support the potential clinical translation of targeting senescence as a supplementary treatment for HFpEF, particularly in the context of CKM syndrome. Several agents have been developed as senolytic agents^35, 36^. Among the most commonly used agents are intermittent treatment with Dasatinib (a tyrosine kinase inhibitor) and Quercetin, as well as Fisetin. Both Quercetin and Fisetin belong to the flavonoid polyphenols, which exhibit antioxidant and anti-inflammatory activities. Indeed, several ongoing clinical trials are evaluating these senolytic agents for various age-related diseases (clinicaltrials.gov). Early phase and pilot clinical trials of senolytics suggest they decrease senescent cells, reduce inflammation, alleviate frailty in humans^37^, and modestly improve SASP in early Alzheimer’s disease^38, 39^. However, given the heterogeneity of senescence and the lack of specificity of currently available senolytic agents, our data using the genetic senolytic approach in p16-3MR provides proof of concept for the future development of agents with greater specificity.

Our p16-3MR mice were designed to address senescence involving any type of cells. Indeed, we observe signals of DNA damage response, senescence and inflammation in multiple cell types in both heart and kidney. However, cell-specific inhibition of cGAS/STING or senescence in the 4-hit mouse model warrants future studies. As our 4-hit mouse model requires long-term experiments, to address the limitation of our current mouse studies, we turned to *Drosophila* to answer cell-type specific inhibition of CGAS/STING pathway. Since no model can perfectly recapitulate human Cardiovascular-Kidney-Metabolic (CKM) syndrome, our cross-species approach showing conservation of these pathways across multiple species provides biological robustness. The translational relevance is further strengthened by using human iPS-CM model and validation in human HFpEF biopsy specimens.

In conclusion, using mouse, fly, and human iPSC models and human cardiac biopsies, we elucidate the critical role of senescence and DNA damage sensors (cGAS/STING and ZBP1) in activating inflammation in response to chronic cardiovascular-kidney-metabolic (CKM) stress, which clinically manifests as HFpEF. Our findings suggest a potential clinical translation of targeting senescence and innate immune pathways as promising new therapeutics for HFpEF, particularly for those with obesity, diabetes, and cardiovascular kidney metabolic syndrome.

## Author Contribution

**Conceptualization:** D.D. and Z.H. designed the study. **Data acquisition and analysis:** D.D, J.Z, M.G., K.W, X.Y, M.T, D.C A.R, A.H.J., R.L, and V.H. **Writing:** D.D, J.Z and Z.H. wrote the paper with input from all authors. **Supervision:** D.D. and Z.H. were involved in the planning and supervision of the work.

---

**Figure S1.**
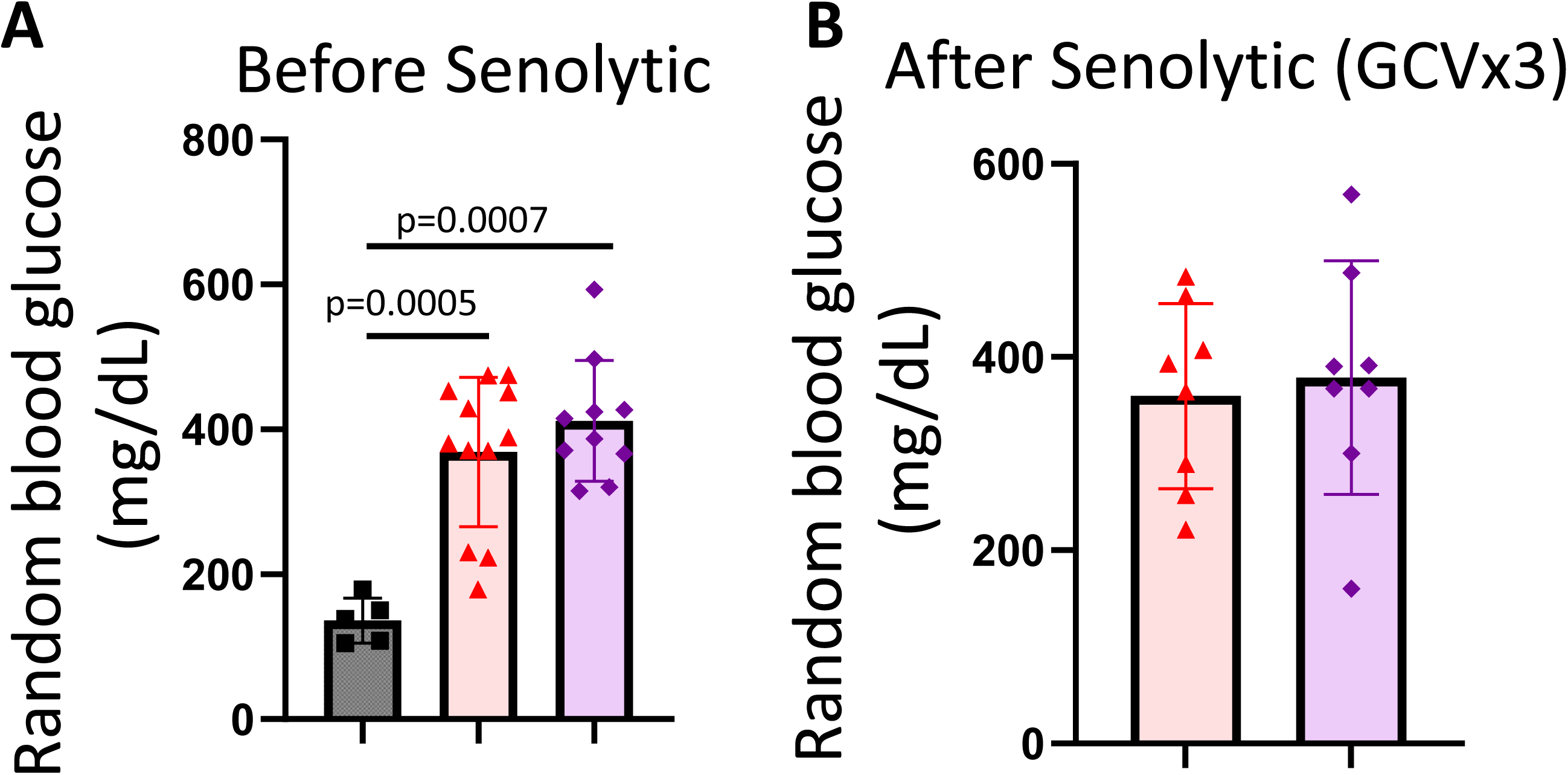
Random blood glucose by OneTouch Glucometer, (A) Before and (B) After Senolytic therapy (3 cycles of ganciclovir).N=5-12; non-parametric tests.

**Figure S2.**
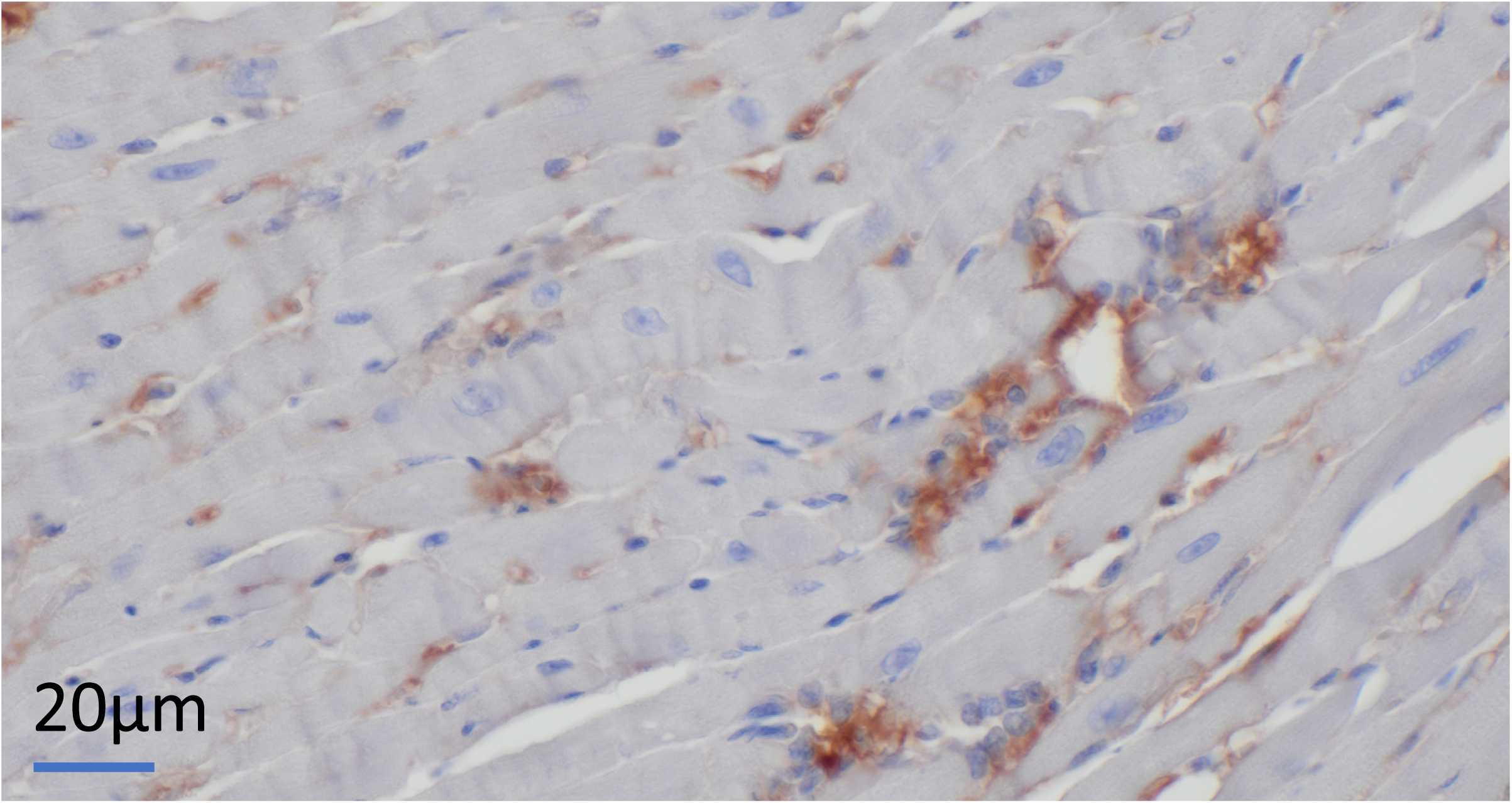
Clusters of CD68-positive cells in 4-Hit mouse heart, matching the clusters of mononuclear inflammatory cells (macrophages) by H&E stains (Figure 2D).

**Figure S3.**
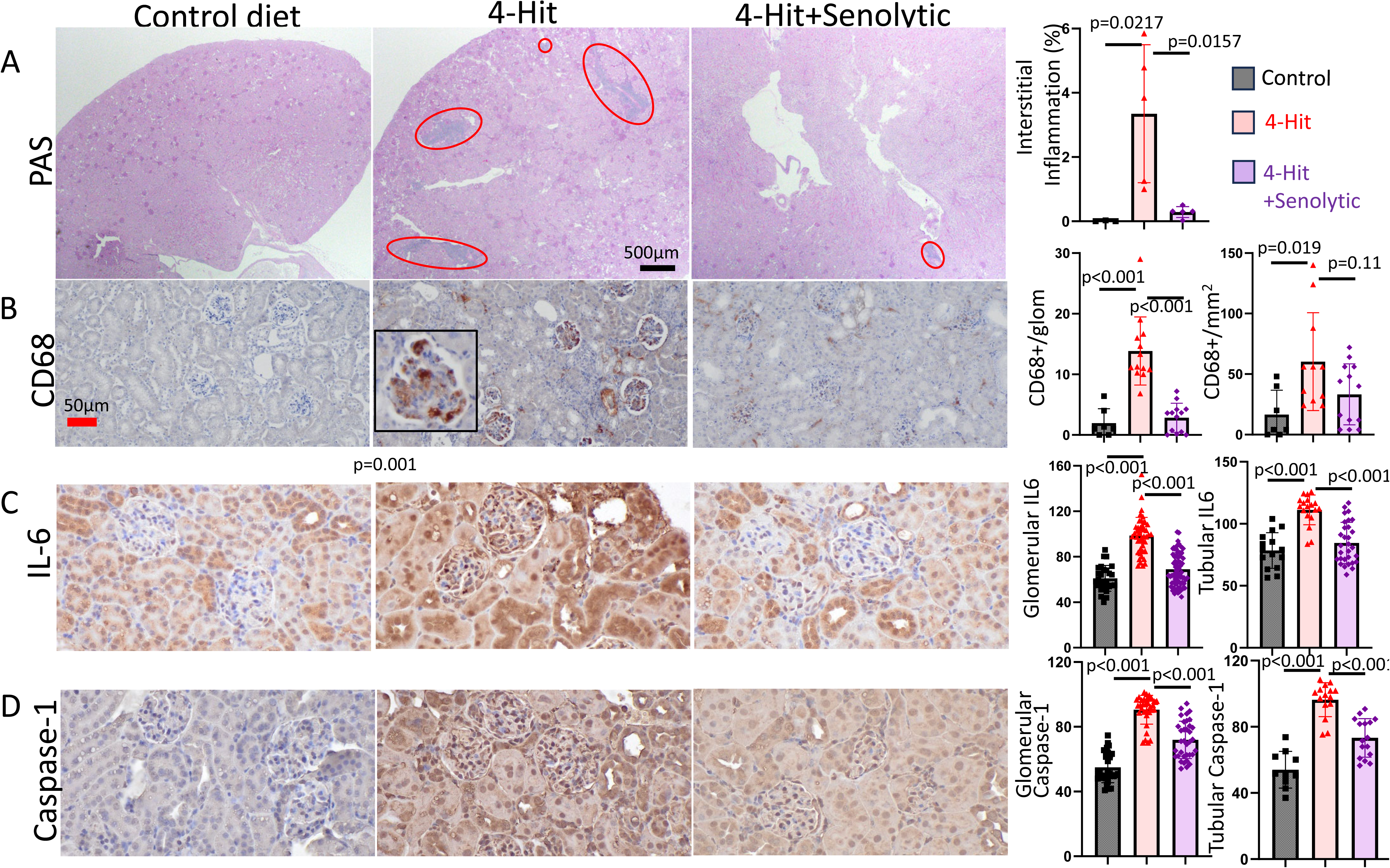
Increased Inflammation in chronic kidney disease (4-Hit CKM) was attenuated by Senolytic treatment. (A). Representative low (2x) magnification of PAS-stained kidneys showing clusters of mononuclear inflammatory infiltrates (red ovals) and the % area of inflammatory cells, (B). IHC of CD68-positive cells within the expanded glomerular mesangium (number of CD68+ cells/glomerulus) and in the interstitium (number of CD68+ cells/ mm^2^). Inset: representative high magnification. (C). IHC of Interleukin 6 (integrated density, arbitrary units) and quantitative analysis for glomeruli and tubulointerstitium. (D). IHC of Caspase-1 (integrated density, arbitrary units) and quantitative analysis for glomeruli and tubulo-interstitium. N=3 control diet mice and 5-6 each group of 4-Hit mice; ANOVA followed by Sidak post-hoc tests.

**Figure S4.**
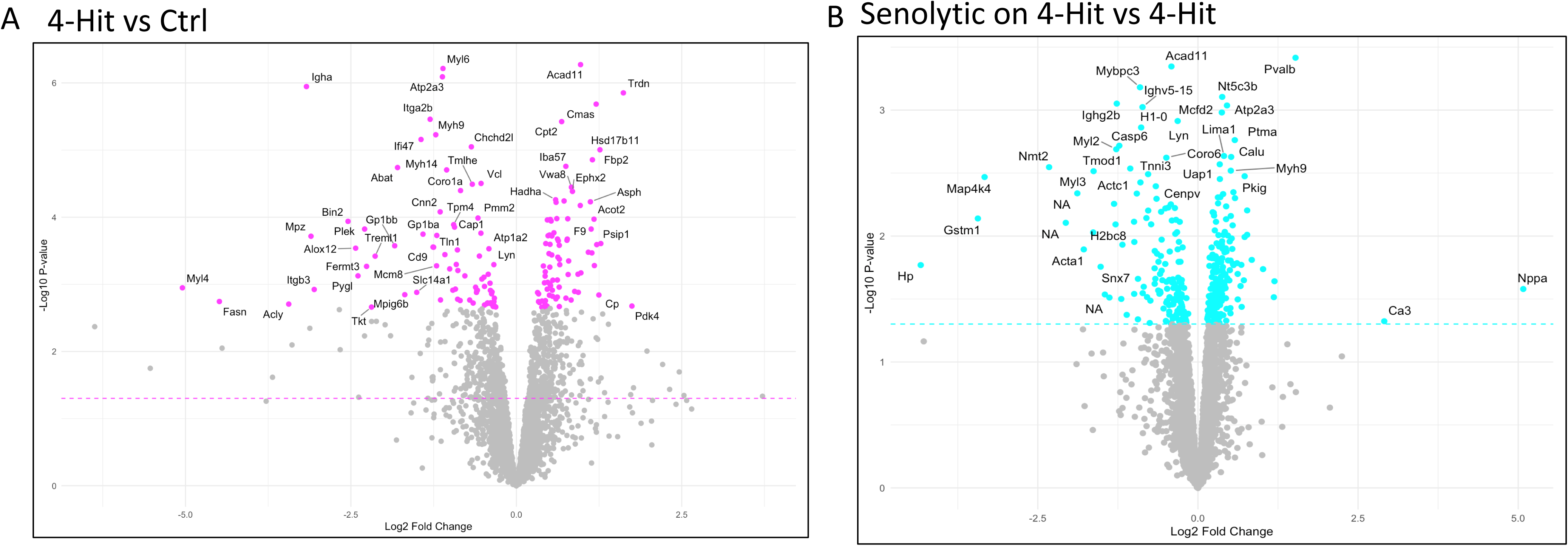
Volcano plots (-Log p-values vs Log2 fold-change) of cardiac proteome changes in 4-Hit mice. (A) 4-Hit vs control diet, showing 4-Hit effect, (B).Senolytic therapy with ganciclovir in p16-3MR mice stressed with 4-Hit vs untreated 4-Hit mouse hearts. Only partial lists of gene names are labeled.

**Figure S5.**
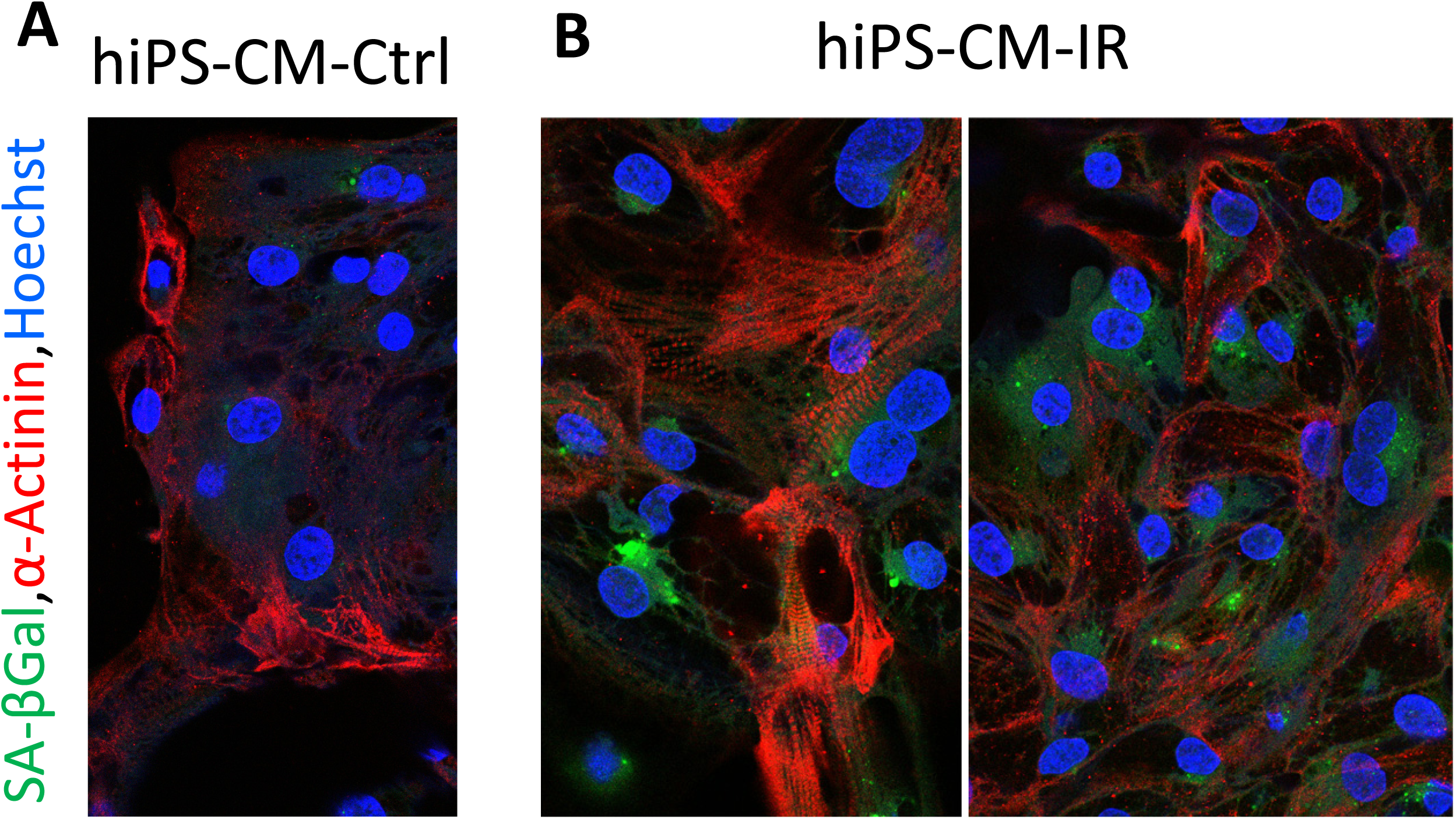
The hiPS-CM were stained with CellEvent™ Senescence Green Assay Kit and α-Actinin cardiomyocyte marker. (A). Control hiPS-CM, (B). hiPS-CM stressed with irradiation, showing green signals of Senescence-Associated β-Galactosidase.

**Figure S6.**
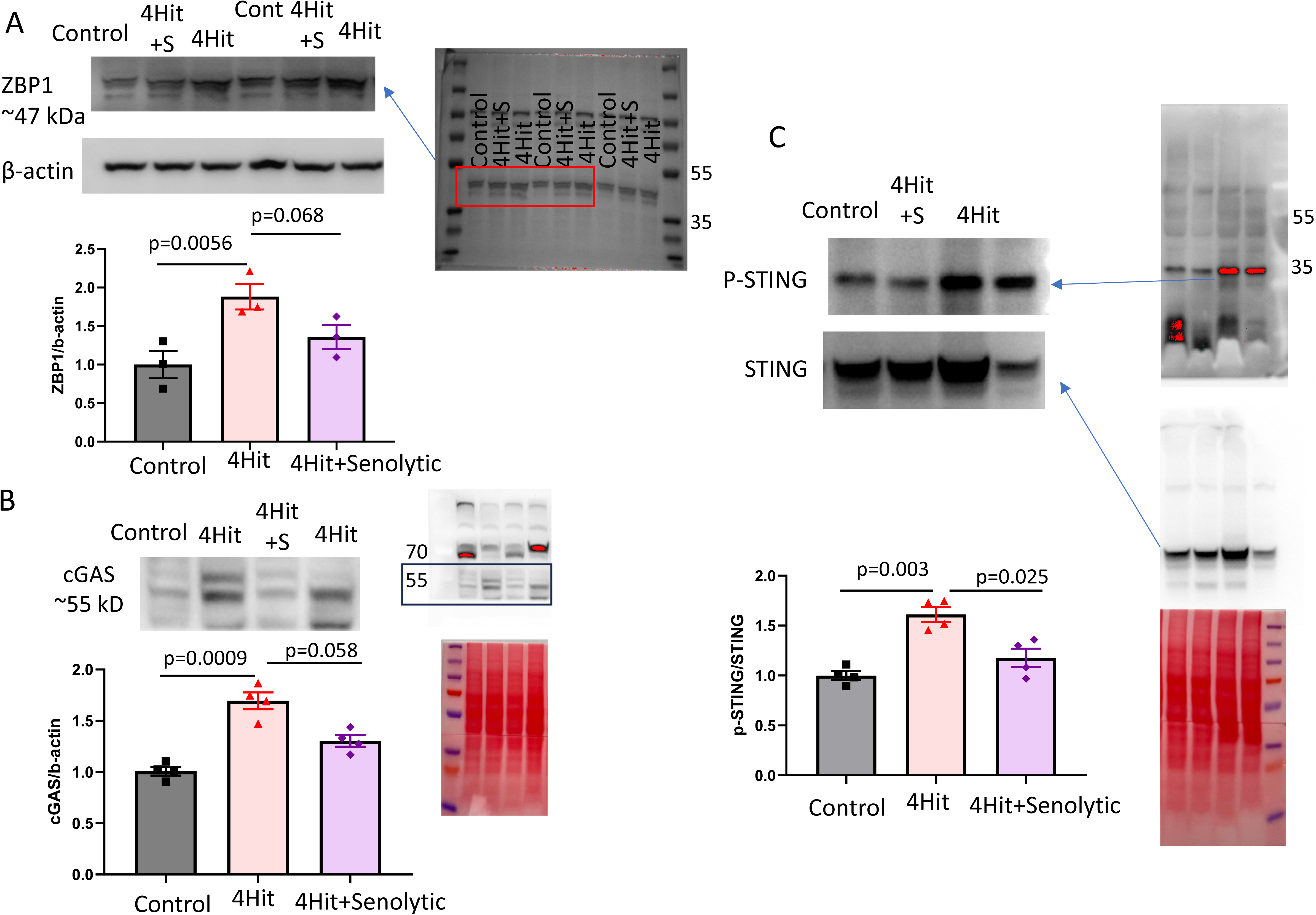
Increased ZBP1/cGAS/STING in 4-Hit CKM mouse hearts was attenuated by Senolytic treatment. (A). ZBP1, (B). cGAS. (C). Phospho and total STING. N=3-4 each group; ANOVA followed by Dunn post-hoc tests.

